## Supplemental Table S1 for "PHFinder: Assisted detection of point heteroplasmy in Sanger sequencing chromatograms"

Supporting Information for **PHFinder**: Assisted detection of point heteroplasmy in Sanger sequencing chromatograms.

**Table S1.** Samples with known heteroplasmies in the dataset, AB1 files, position of the heteroplasmy in the reference sequence (NC_006927.1) and MR at the heteroplasmic nucleotide site.

| Sample ID | AB1 file | Position | Main ratio |
|---|---|---|---|
| MN001 | 5092_6.ab1 | 258 | 88 |
| MN001 | 5095_14.ab1 | 258 | 73 |
| MN001 | 5097_4.ab1 | 258 | 76 |
| MN010 | 5985_38.ab1 | 258 | 40 |
| MN014 | 5982_53.ab1 | 143 | 58 |
| MN020 | 5981_17.ab1 | 143 | 21 |
| MN026 | 5500_3.ab1 | 82 | 32 |
| MN027 | 5497_38.ab1 | 82 | 86 |
| MN043 | 5504_9.ab1 | 235 | 63 |
| MN033 | 5515_54.ab1 | 82 | 24 |
| MN035 | 5512_12.ab1 | 258 | 68 |
| MN013 | 5507_2.ab1 | 82 | 41 |
| MN056 | 4731_18.ab1 | 281 | 53 |
| MN060 | 4751_17.ab1 | 258 | 47 |
| MN097 | 4751_15.ab1 | 282 | 59 |
| MN097 | 4744_38.ab1 | 282 | 46 |
| MN061 | 4744_32.ab1 | 82 | 21 |
| MN062 | 4729_37.ab1 | 143 | 26 |
| MN063 | 4661_65.ab1 | 281 | 75 |
| MN063 | 4666_19.ab1 | 281 | 76 |
| MN063 | 5101_11.ab1 | 281 | 78 |
| MN064 | 4661_53.ab1 | 258 | 29 |
| MN065 | 4734_3.ab1 | 143 | 21 |
| MN066 | 4661_37.ab1 | 82 | 31 |
| MN066 | 5101_3.ab1 | 82 | 33 |
| MN069 | 4661_8.ab1 | 281 | 46 |

| | | | |
|---|---|---|---|
| MN071 | 4661_1.ab1 | 258 | 49 |
| MN071 | 4666_17.ab1 | 258 | 58 |
| MN058 | 4655_6.ab1 | 258 | 60 |
| MN058 | 4667_80.ab1 | 258 | 59 |
| MN081 | 5509_19.ab1 | 258 | 73 |
| MN085 | 4948_82.ab1 | 82 | 29 |
| MN086 | 4948_79.ab1 | 143 | 73 |
| MN086 | 5311_26.ab1 | 143 | 64 |
| MN088 | 4948_62.ab1 | 143 | 27 |
| MN088 | 5138_89.ab1 | 143 | 20 |
| MN090 | 4626_23.ab1 | 258 | 70 |
| MN095 | 5138_28.ab1 | 258 | 41 |
| MN095 | 4954_29.ab1 | 258 | 74 |
| MN017 | 4598_77.ab1 | 235 | 27 |
| MN017 | 4667_38.ab1 | 235 | 18 |
| MN098 | 4598_53.ab1 | 258 | 48 |
| MN098 | 4667_36.ab1 | 258 | 40 |

Notes: Positions based on the region analysed; position 1 corresponds to position 15,490 in the reference NC_006927.1 (NCBI Reference Sequence) from Sasaki et al. (2005).

**Table S2.** Summary of the results for each of the 64 sets of index threshold values.

| Parameters | HP_AB1 | HP_Samples | False_Positives | Analysed_Samples | Analysed_AB1 | Open_AB1 |
|---|---|---|---|---|---|---|
| MR15-SR0.2-AQ30 | 21 | 16 | 1 | 100 | 143 | 22 |
| MR15-SR0.2-AQ40 | 20 | 15 | 1 | 88 | 115 | 21 |
| MR15-SR0.2-AQ50 | 14 | 12 | 0 | 68 | 79 | 14 |
| MR15-SR0.2-AQ60 | 3 | 3 | 0 | 37 | 15 | 3 |
| MR15-SR0.3-AQ30 | 31 | 22 | 9 | 100 | 143 | 38 |
| MR15-SR0.3-AQ40 | 30 | 21 | 5 | 88 | 115 | 34 |
| MR15-SR0.3-AQ50 | 21 | 15 | 2 | 68 | 79 | 23 |
| MR15-SR0.3-AQ60 | 4 | 4 | 0 | 37 | 15 | 4 |
| MR15-SR0.4-AQ30 | 39 | 28 | 38 | 100 | 143 | 66 |
| MR15-SR0.4-AQ40 | 38 | 27 | 23 | 88 | 115 | 52 |
| MR15-SR0.4-AQ50 | 29 | 21 | 13 | 68 | 79 | 36 |
| MR15-SR0.4-AQ60 | 7 | 7 | 0 | 37 | 15 | 7 |
| MR15-SR0.5-AQ30 | 41 | 30 | 113 | 100 | 143 | 94 |
| MR15-SR0.5-AQ40 | 39 | 28 | 61 | 88 | 115 | 67 |
| MR15-SR0.5-AQ50 | 30 | 22 | 25 | 68 | 79 | 42 |
| MR15-SR0.5-AQ60 | 7 | 7 | 2 | 37 | 15 | 8 |
| MR20-SR0.2-AQ30 | 21 | 16 | 1 | 100 | 143 | 22 |
| MR20-SR0.2-AQ40 | 20 | 15 | 1 | 88 | 115 | 21 |
| MR20-SR0.2-AQ50 | 14 | 12 | 0 | 68 | 79 | 14 |
| MR20-SR0.2-AQ60 | 3 | 3 | 0 | 37 | 15 | 3 |
| MR20-SR0.3-AQ30 | 30 | 22 | 7 | 100 | 143 | 35 |
| MR20-SR0.3-AQ40 | 29 | 21 | 4 | 88 | 115 | 32 |
| MR20-SR0.3-AQ50 | 20 | 15 | 1 | 68 | 79 | 21 |
| MR20-SR0.3-AQ60 | 4 | 4 | 0 | 37 | 15 | 4 |
| MR20-SR0.4-AQ30 | 38 | 28 | 18 | 100 | 143 | 52 |
| MR20-SR0.4-AQ40 | 37 | 27 | 8 | 88 | 115 | 43 |
| MR20-SR0.4-AQ50 | 28 | 21 | 4 | 68 | 79 | 32 |
| MR20-SR0.4-AQ60 | 7 | 7 | 0 | 37 | 15 | 7 |
| MR20-SR0.5-AQ30 | 40 | 30 | 66 | 100 | 143 | 78 |
| MR20-SR0.5-AQ40 | 38 | 28 | 31 | 88 | 115 | 57 |
| MR20-SR0.5-AQ50 | 29 | 22 | 11 | 68 | 79 | 37 |
| MR20-SR0.5-AQ60 | 7 | 7 | 1 | 37 | 15 | 8 |
| MR25-SR0.2-AQ30 | 20 | 15 | 1 | 100 | 143 | 21 |
| MR25-SR0.2-AQ40 | 19 | 14 | 1 | 88 | 115 | 20 |
| MR25-SR0.2-AQ50 | 13 | 11 | 0 | 68 | 79 | 13 |
| MR25-SR0.2-AQ60 | 3 | 3 | 0 | 37 | 15 | 3 |
| MR25-SR0.3-AQ30 | 28 | 21 | 5 | 100 | 143 | 31 |
| MR25-SR0.3-AQ40 | 27 | 20 | 3 | 88 | 115 | 29 |
| MR25-SR0.3-AQ50 | 18 | 14 | 0 | 68 | 79 | 18 |

| | | | | | | |
|---|---|---|---|---|---|---|
| MR25-SR0.3-AQ60 | 3 | 3 | 0 | 37 | 15 | 3 |
| MR25-SR0.4-AQ30 | 34 | 25 | 9 | 100 | 143 | 40 |
| MR25-SR0.4-AQ40 | 33 | 24 | 4 | 88 | 115 | 35 |
| MR25-SR0.4-AQ50 | 24 | 18 | 0 | 68 | 79 | 24 |
| MR25-SR0.4-AQ60 | 5 | 5 | 0 | 37 | 15 | 5 |
| MR25-SR0.5-AQ30 | 35 | 26 | 40 | 100 | 143 | 57 |
| MR25-SR0.5-AQ40 | 34 | 25 | 18 | 88 | 115 | 44 |
| MR25-SR0.5-AQ50 | 25 | 19 | 6 | 68 | 79 | 30 |
| MR25-SR0.5-AQ60 | 5 | 5 | 1 | 37 | 15 | 6 |
| MR30-SR0.2-AQ30 | 19 | 14 | 1 | 100 | 143 | 20 |
| MR30-SR0.2-AQ40 | 18 | 13 | 1 | 88 | 115 | 19 |
| MR30-SR0.2-AQ50 | 12 | 10 | 0 | 68 | 79 | 12 |
| MR30-SR0.2-AQ60 | 2 | 2 | 0 | 37 | 15 | 2 |
| MR30-SR0.3-AQ30 | 25 | 18 | 3 | 100 | 143 | 28 |
| MR30-SR0.3-AQ40 | 24 | 17 | 2 | 88 | 115 | 26 |
| MR30-SR0.3-AQ50 | 15 | 11 | 0 | 68 | 79 | 15 |
| MR30-SR0.3-AQ60 | 2 | 2 | 0 | 37 | 15 | 2 |
| MR30-SR0.4-AQ30 | 29 | 20 | 5 | 100 | 143 | 33 |
| MR30-SR0.4-AQ40 | 28 | 19 | 3 | 88 | 115 | 30 |
| MR30-SR0.4-AQ50 | 19 | 13 | 0 | 68 | 79 | 19 |
| MR30-SR0.4-AQ60 | 3 | 3 | 0 | 37 | 15 | 3 |
| MR30-SR0.5-AQ30 | 30 | 21 | 25 | 100 | 143 | 46 |
| MR30-SR0.5-AQ40 | 29 | 20 | 15 | 88 | 115 | 37 |
| MR30-SR0.5-AQ50 | 20 | 14 | 4 | 68 | 79 | 23 |
| MR30-SR0.5-AQ60 | 3 | 3 | 0 | 37 | 15 | 3 |

Notes: HP_AB1: Number of AB1 files with an heteroplasmy detected. HP_Samples: Number of heteroplasmic samples detected. False_Positives: Number of false positives detected. Analysed_Samples: Number of samples analysed by PHFinder. Open_AB1: Number of AB1 that would need to be manually checked.
